# Changes in historical typhoid transmission across 16 U.S. cities, 1889-1931: Quantifying the impact of investments in water and sewer infrastructures

**DOI:** 10.1101/609677

**Authors:** Maile T. Phillips, Katharine A. Owers, Bryan T. Grenfell, Virginia E. Pitzer

## Abstract

**Background:** Investments in water and sanitation systems are believed to have led to the decline in typhoid fever in developed countries, such that most cases now occur in regions lacking adequate clean water and sanitation. Exploring seasonal and long-term patterns in historical typhoid mortality in the United States can offer deeper understanding of disease drivers.

**Methods:** We fit modified Time-series Susceptible-Infectious-Recovered models to city-level weekly mortality counts to estimate seasonal and long-term typhoid transmission. We examined seasonal transmission separately by city and aggregated by water source. We fit regression models to measure associations between long-term transmission and financial investments in water and sewer systems.

**Results:** Typhoid transmission peaked in late summer/early fall. Seasonality varied by water source, with the greatest variation occurring in cities with reservoirs. Historical $1 per capita ($25.80 in 2017) investments in construction and operation of water and sewer systems were associated with 8-53% decreases in typhoid transmission, while $1 increases in total value or debt accrued to maintain them were associated with 4-7% decreases.

**Conclusion:** Our findings aid in the understanding of typhoid transmission dynamics and potential impacts of water and sanitation improvements, and can inform cost-effectiveness analyses of interventions to reduce the typhoid burden.

## INTRODUCTION

Typhoid fever is caused by infection with the bacteria *Salmonella enterica* serovar Typhi, which is mainly transmitted through faecal contamination of food or water (1). In many developed countries, including the United States (U.S.), investments in water and sewer infrastructures led to the decline in typhoid incidence in the beginning of the 20^th^ century, such that the majority of the global burden now occurs in countries where sanitary conditions are poor and access to clean water and sanitation is lacking (1–4).

Examining short- and long-term trends in typhoid incidence can provide insights into factors driving transmission (5). In many countries, typhoid fever follows a seasonal pattern, with peak incidence occurring around the same time every year (6). Seasonality in typhoid exhibits distinct patterns by region and latitude, and can be influenced by rainfall, temperature, and climate (7). However, drivers of seasonal patterns in typhoid are not yet fully understood.

Long-term patterns in typhoid cases have also been investigated, particularly in countries where cases have declined to almost zero (5). In the U.S., the number of typhoid cases decreased from a reported 35,000 in 1900 to fewer than 300 in 2017 despite a 4.3-fold population increase (8–11). While it is commonly accepted that investments in water and sanitation are responsible for the decline in typhoid fever, there is limited empirical evidence to support this claim. In one study, Cutler and Miller found that the introduction of clean water technologies was responsible for almost half of the mortality reduction in major cities at the beginning of the 20^th^ century; however, they did not consider complexities of the transmission process (12).

In this study, we developed a mathematical model of typhoid transmission to examine seasonal and long-term trends in typhoid transmission from 1889-1931 in 16 U.S. cities. Our objectives were two-fold: (1) to examine how seasonal patterns of typhoid transmission varied geographically and historically depending on the water supply and treatment; and (2) to quantify the relationship between investments in water and sanitation infrastructures and long-term typhoid transmission rates.

## METHODS

### Study Design, Data, and Variables

We extracted reported weekly typhoid mortality from 1889 to 1931 at the city level from the Project Tycho database (13, 14). Cities were chosen based on two criteria: (1) at least 1,000 typhoid deaths were reported during the study period, and (2) less than 25% of weekly data was missing. These exclusion criteria resulted in data for 16 U.S. cities (Figure S1).

Yearly population estimates were obtained from the U.S. Census Bureau (15–17). The population <1 year of age was used as a proxy for births, since birth rate data was not available and typhoid is rare in <1-year-olds (18). Cubic splines were used to extrapolate weekly population estimates. Financial data on water supply and sewer systems for each city were extracted from U.S. Census Bureau yearly reports (16); we obtained data on eight variables across five categories: “receipts,” “expenses,” “outlays,” “total value,” and “funded debt” (Table 1), divided by the yearly city population to generate per capita estimates. Data on water treatment and type for each city were extracted from a variety of sources (Table S1) (17).

**Table 1.** Definitions of financial variables. Each of the five categories of financial variables used in this study are described, as defined by the U.S. Census Bureau in its annual “Financial Statistics” series (the source of these variables).

| Description |  |
| --- | --- |
| **Maintenance and operation** |  |
| Receipts | Receipts for payments for governmental costs. These receipts usually take the form of money, bills receivable, land, and services. All city revenue receipts were recorded in the city books for municipally-operated water supply and sewer systems for the public or city (excluding interest from current deposits). |
| Expenses | City government costs, other than interest, of (1) services employed, property rented, and materials consumed in connection with maintenance and operation; (2) losses from deflation, bank failures, and related causes; and (3) depreciation of permanent properties and public improvements. |
| **Acquisition and construction** |  |
| Outlays | Total annual amounts paid by the city for the acquisition or construction of permanent lands, properties and public improvements. These include payments for additions made to previously acquired or constructed properties. |
| **Overall value and debt** |  |
| Value | Total estimated value of the public properties (including depreciation), including both the business value and the physical value of the building and equipment. This amount is estimated separately by city officials, and is acknowledged to not be estimated uniformly across cities. |
| Funded debt | Long-term debts or debt liabilities in the form of bonds or certificates of indebtedness that the city government is under obligation to pay. |

All cities had missing data on weekly typhoid mortality. In many cases, missing mortality counts were instances of zero cases, because cities frequently only reported during weeks when deaths occurred. To account for both true zero counts and missing data, mortality data were coded as zeroes if there were fewer than 13 consecutive weeks of missing death counts, and imputed as missing data if there were 13 or more consecutive weeks (using “imputeTS” (19)). We ran a sensitivity analysis to assess how this arbitrary 13-week cut-off could impact our results. After imputation, weekly typhoid mortality counts and population estimates were aggregated into four-week periods to approximate the generation interval of typhoid (20). Since the mortality data were later log-transformed, we added one to every four-week data point; a sensitivity analysis was again performed to assess the impact of adding different values.

### Statistical Methods

We conducted preliminary analyses to describe differences in typhoid mortality trends between cities and pre- to post-intervention. First, we fit generalized linear models (GLMs) with linear time trends and one-year and six-month harmonics to the pre- and post-intervention time series (defined as two years after the first water supply intervention) for each city. We compared intercepts, slopes, and six-month and one-year amplitudes for the pre- and post- periods.

We then fit Time-series Susceptible-Infectious-Recovered (TSIR) models (21) to each city’s pre- and post-intervention time series to investigate seasonal and long-terms trends in typhoid transmission rates. Briefly, TSIR models estimate the disease transmission rate by reconstructing the underlying susceptible and infectious populations. New infections at time *t*+1 (*I*_*t*+1_) arise from transmission from infectious (*I*_*t*_) to susceptible (*S*_*t*_) individuals at time *t*:

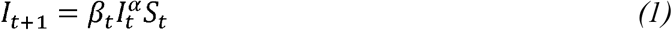

where *β*_*t*_ is the disease transmission rate at time *t*. The exponent *α* allows for heterogeneous population mixing and corrects for discretization of the continuous-time infection process (22).

We reconstructed the infectious (*I*_*t*_) and susceptible (*S*_*t*_) populations using the first ten years of typhoid mortality and census data (prior to the introduction of water and sanitation interventions) (21, 23). The susceptible population at time *t* is equal to the previous susceptible population plus new births minus new infections, summarized as follows:

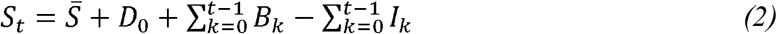

where *S̅* is the mean susceptible population, *D*_0_ is the deviation at time zero, 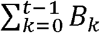 is the sum of births up to time *t*, and 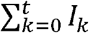 is the sum of infections up to time *t*. The number of “true” infections at time *t* is estimated from the observed deaths at time *t* divided by an underreporting factor *ρ*(*I*_*t*_ = *Y*_*t*_/*ρ*), which also accounts for the case fatality rate. Equation 2 can be rearranged as

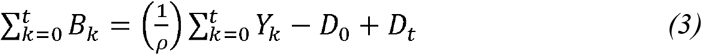

to estimate the underreporting fraction, deviation at time zero, and model residuals (*D*_*t*_ = *S*_*t*_ − *S̅*).

We modified Equation 1 to account for the unique features of typhoid epidemiology, including the contribution of chronic carriers (*C*) to the prevalence of infection. Furthermore, we separated the transmission parameter *β*_*t*_ into seasonal and long-term components (*β*_*seas*_ and *β*_*lt*_, respectively). Thus, the TSIR model for typhoid is as follows:

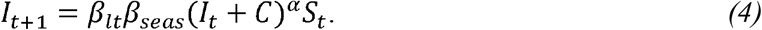

Equation 4 can be log-transformed:

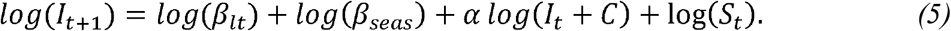

To estimate *C* and *S*_*t*_ (= *D*_*t*_ = *S̅*), we maximized the likelihood of the fitted regression over different values of *C* and *S̅*, each ranging from 0 to the population size. We then used the best-fit model to estimate seasonal and long-term typhoid transmission rates.

For our main analysis, we further modified the TSIR model (Equation 5) to include waning of immunity. We modelled the susceptible population at time *t* as a function of the total population at time *t* minus the previously infectious and recovered individuals: 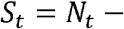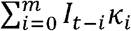, where *κ*_*i*_ is the degree of immunity *i* generation intervals after infection. These methods are described in detail elsewhere (21, 24, 25); the model fitting and validation process is described in the Supplemental Material.

### Examining predictors of seasonal and long-terms trends in transmission

Once we identified the optimal model for each city, we extracted the seasonal and long-term transmission rates. Seasonal transmission parameters were plotted separately for each city and aggregated by water source type.

To examine associations between long-term typhoid transmission and financial investments in water and sewer systems, we fit linear regression models for each city and financial variable separately. The yearly-average log-transformed long-term transmission parameter was the outcome and each financial variable was a univariate predictor. Complete case analysis was assumed for missing financial variable data.

All analyses were performed using R version 1.0.153 (26).

## RESULTS

From 1889-1931, there were 86,023 typhoid deaths across all cities (median: 3,382 deaths per city). Figure S2 shows the weekly time series of typhoid mortality in each city. Of the 16 cities, four used reservoirs or lakes as their water source, three drew water from the Great Lakes, and nine accessed water from rivers (Table 2). Most cities introduced water chlorination or filtration during the study period, but some cities implemented other interventions. Boston’s Metropolitan Water District completed a new reservoir in 1908, while New York built several additional reservoirs between 1905-1915. The Sanitary District of Chicago changed the direction of flow of the Chicago River so sewage from the city would no longer be discharged into Lake Michigan, the city’s water supply. To address flooding problems from periodic hurricanes and its location below sea level, the New Orleans Drainage Commission began to periodically drain the water supply in 1900. San Francisco had no water supply interventions that we could identify; however, a major earthquake in 1906 resulted in severe infrastructure damage and changes to the water supply system, and was included as a proxy intervention in our analysis.

**Table 2.**
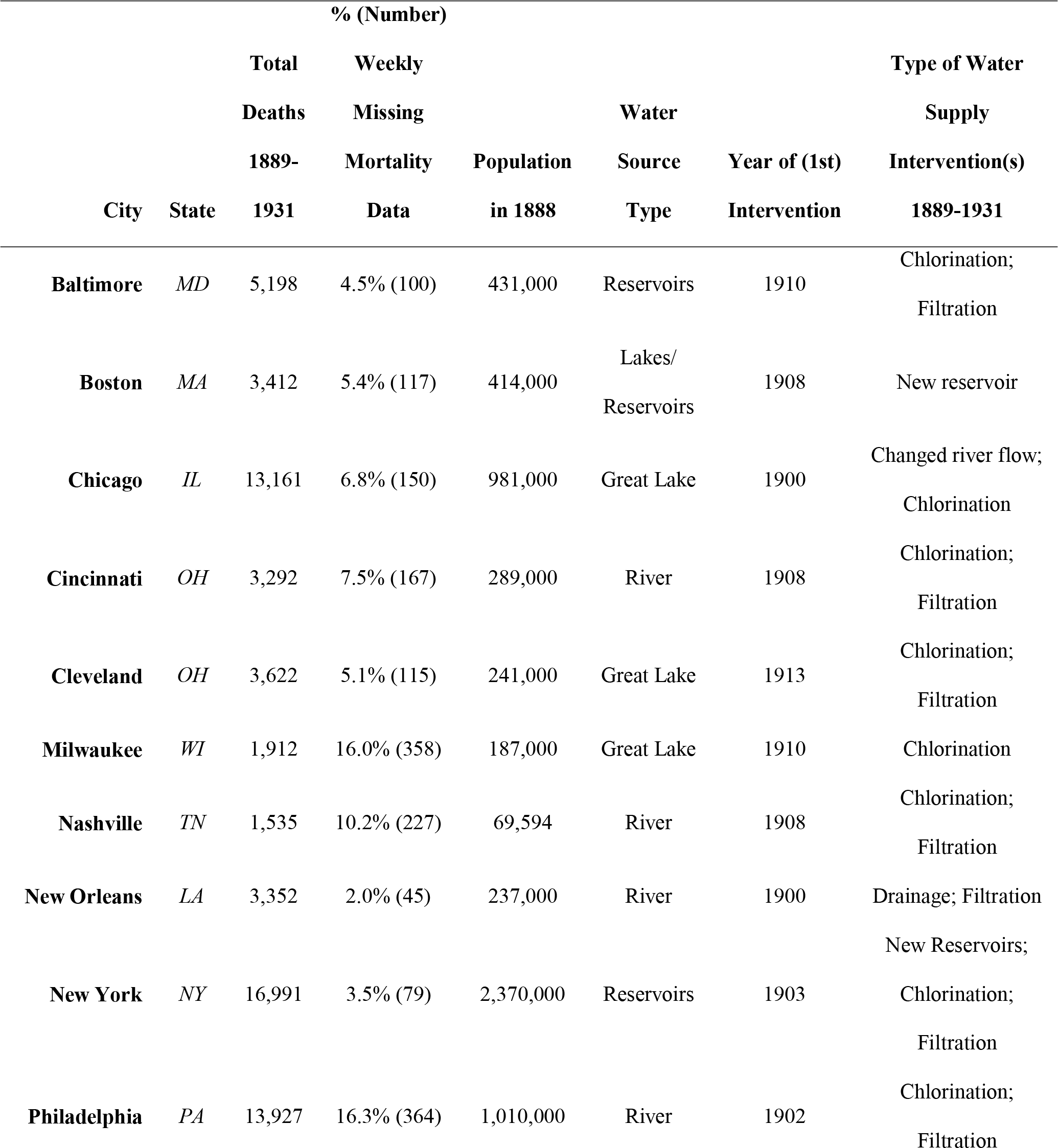
Descriptive statistics of cities and their water supplies. “Total Deaths” are the number reported after imputation for missing data. Missing data numbers represent estimates after correcting for “true zeros” in the datasets, and before imputation.

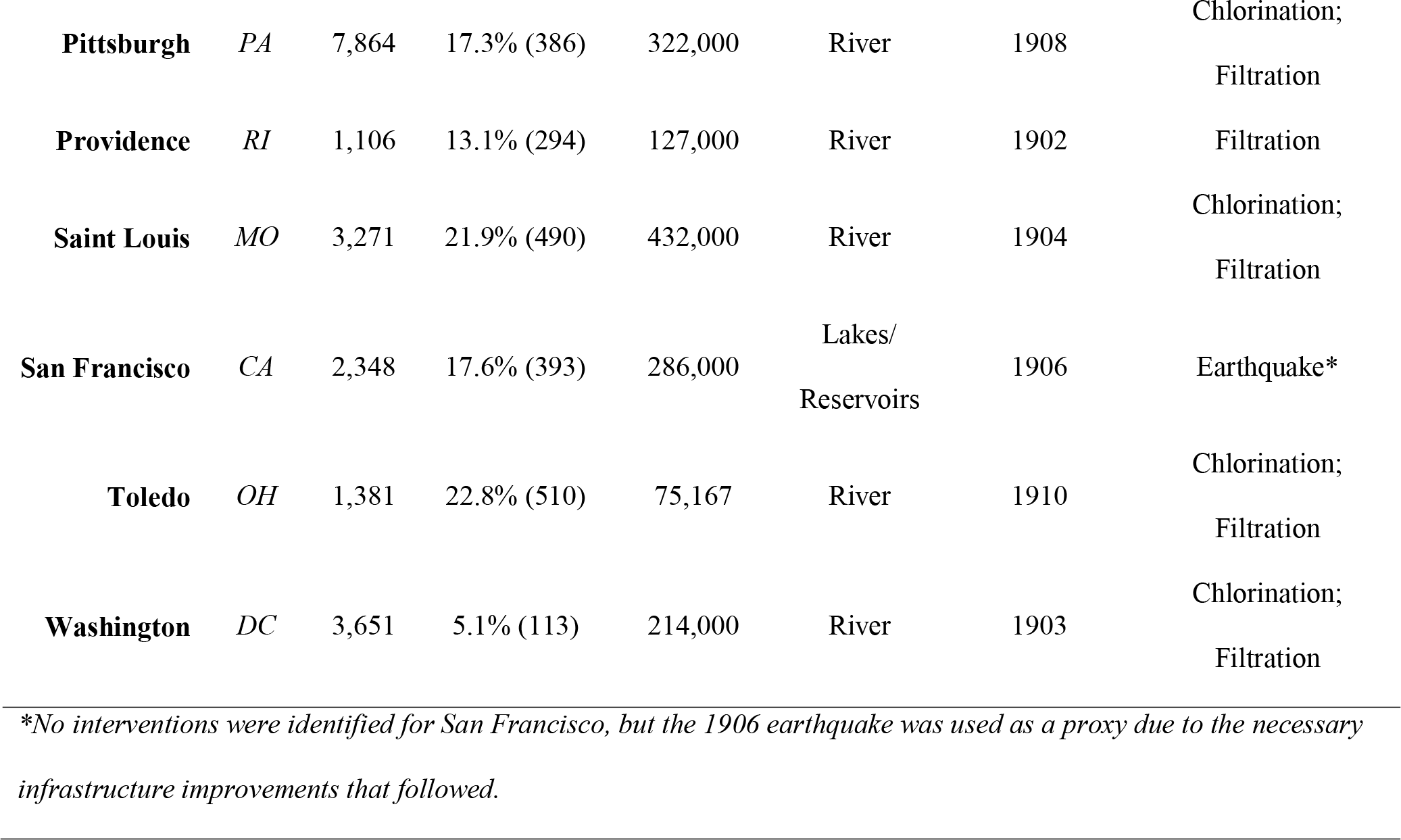

In the preliminary harmonic regression analysis, the six-month amplitude in typhoid mortality decreased in all but two cities (Milwaukee and Nashville), while the one-year seasonal amplitude decreased in all cities but New Orleans post-intervention (Table S2). In every city, typhoid mortality significantly decreased with time in the post-intervention period. The pre-intervention time trend was less consistent across cities.

While the harmonic regression analyses suggested changes in the seasonality of typhoid mortality following interventions, there was little to no difference in seasonality of typhoid transmission pre-versus post-intervention estimated using TSIR models (Figure S3). Thus, we estimated the seasonal transmission rate for the entire 43-year study period in subsequent analyses.

Based on the full TSIR model (including waning of immunity), in most cities typhoid transmission increased at the beginning of the year and peaked around months 8-10 (Figure 1). This trend varied somewhat across cities. In New Orleans, peak transmission occurred earlier (months 6-7), while in San Francisco the peak occurred later (months 10-11). In several cities, there were additional peaks in the winter (months 1-3).

**Figure 1.**
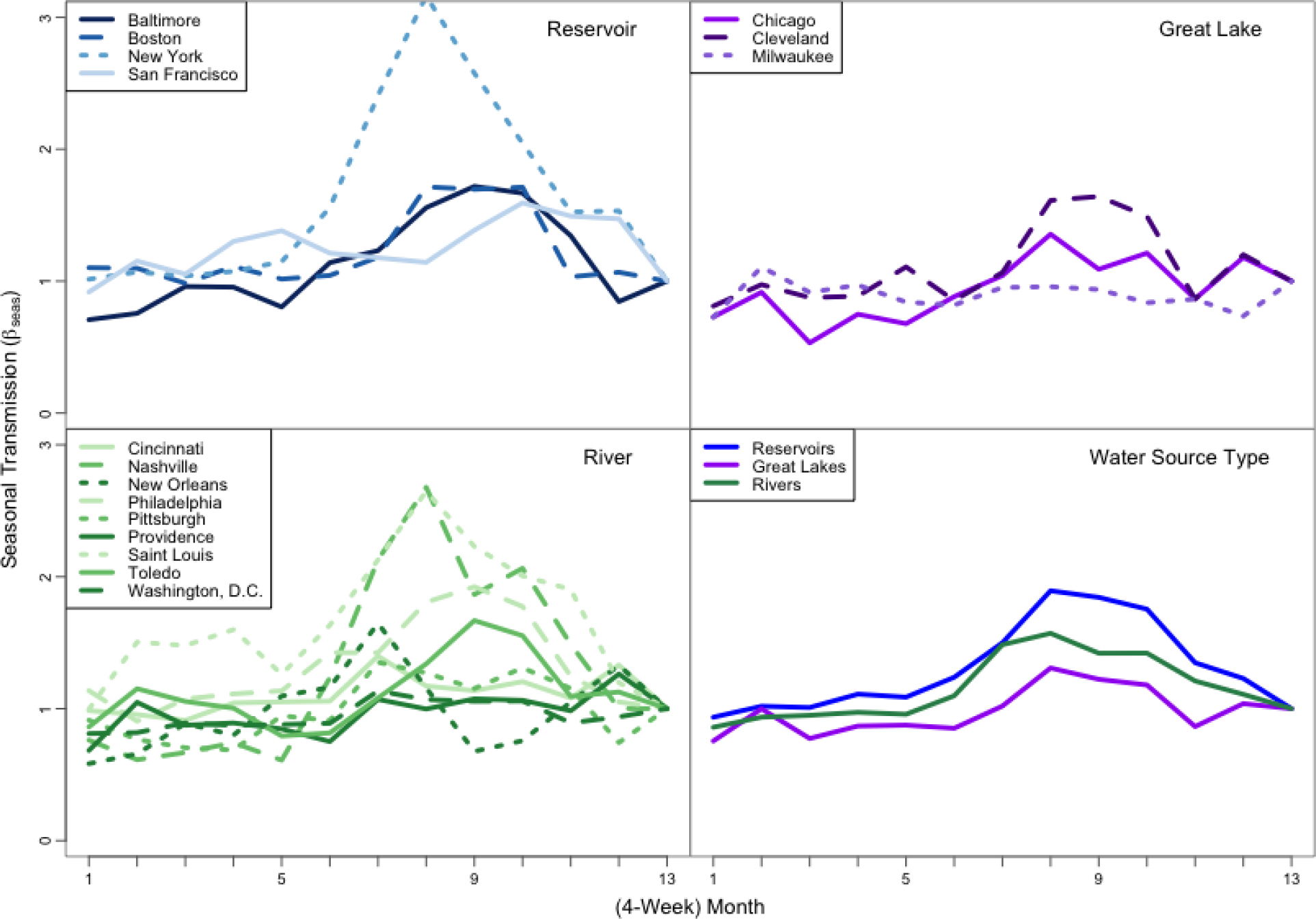
Annual seasonal typhoid transmission estimated from Time-series Susceptible-Infectious-Recovered models. The estimated seasonal transmission rate in each 4-week period is plotted for each city (color-coded by water source type). The last panel shows the seasonal transmission rate by water source type, with reservoirs in blue, rivers in green, and Great Lakes in purple.

Seasonality in typhoid transmission also varied by water source type. While the seasonal trend was similar across water source type, the magnitude of the peaks in transmission differed (Figure 1). Cities that relied on reservoirs had the highest amplitude of seasonal typhoid transmission, while cities that drew water from the Great Lakes had the least variability.

After the 1900s, long-term typhoid transmission began to decrease almost monotonically in every city (Figure 2). In some cities, the decrease in typhoid transmission preceded the first reported water supply intervention, while in others it coincided with or followed the introduction of water supply interventions.

**Figure 2.**
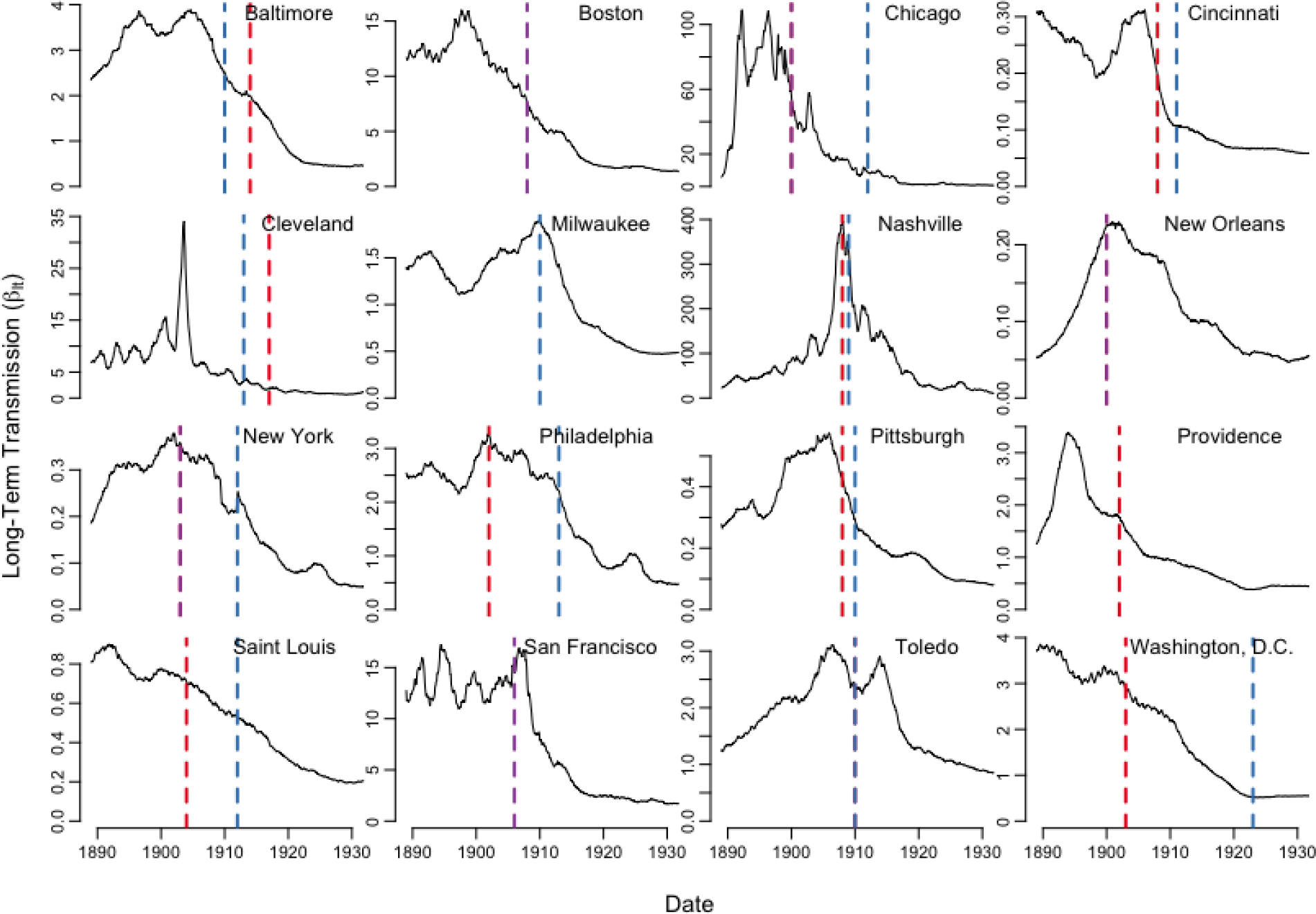
Long-term typhoid transmission by city estimated from Time-series Susceptible-Infectious-Recovered models. The estimated long-term transmission rate (*β*_*lt*_) is plotted in black for each city, while the year in which interventions were introduced are represented by the dashed lines for filtration (red), chlorination (blue), or other interventions (purple).

Investments in both water and sewer systems generally increased over time (Figures S4-S11). Most of the variables related to investments in the water supply were significantly associated with long-term typhoid transmission. Each $1 per capita ($25.80 in 2017 dollars) increase in water supply receipts or expenses was associated with an approximately 45-53% decrease in the typhoid transmission rate, while each $1 per capita increase in the value or funded debt of the water supply was associated with an approximately 4-6% decrease (Table 3). Water supply outlays were associated with more modest decreases in typhoid transmission, likely due to the one-time introduction of water supply interventions. In Cincinnati and Pittsburgh, increases in water supply outlays were associated with increased typhoid transmission. In these two cities, there were large one-time investments in the water supply at the beginning of the study period, which were followed by smaller outlays and decreases in typhoid transmission (Figure S7).

**Table 3.**
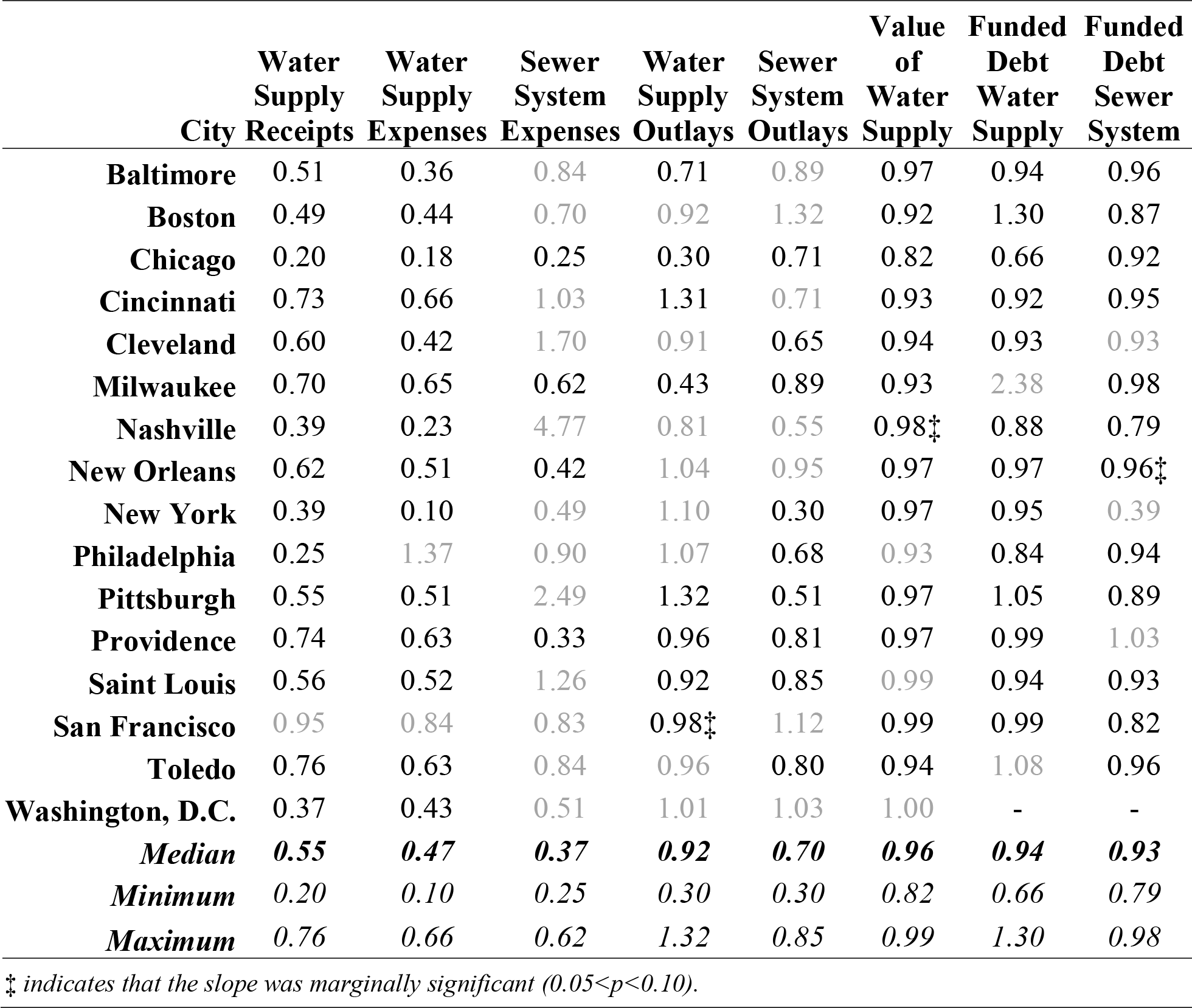
Results of regression analyses: Yearly average long-term typhoid transmission vs. financial investments in water and sewer systems. Each estimate shows the associated change in typhoid transmission for each $1 per capita increase in the financial variable. No data were available for Washington, D.C. for the variables *Funded Debt of the Water Supply System* or *Funded Debt of the Sewer System*. The minimum and maximum values shown in each column are calculated only among cities for which the variable was significant. Black text indicates that the slope had p<0.05. Grey values indicate that the slope was not statistically significant.

Investments in the sewer system were not as strongly associated with long-term trends in typhoid transmission. Sewer system expenses were only significantly associated with long-term typhoid transmission in four cities (Table 3). This variable consistently had two outlier years (1904 and 1909), with increased spending that weakened the overall trend. Sewer system outlays, on the other hand, were associated with typhoid transmission in all but seven cities (30% median decrease per $1 per capita spending). A $1 per capita increase in funded debt and loans for the sewer system was associated with an approximately 7% decrease (range: 2-21% decrease) in long-term transmission rate of typhoid, and was significantly associated with typhoid transmission in all but three cities.

Washington, D.C. and San Francisco had few variables that were significantly associated with long-term typhoid transmission rates. Washington, D.C only had data available for six of the eight variables, and only had significant associations for two of those. San Francisco only had significant associations for four of the eight financial variables; it is possible that investments following the 1906 earthquake and fire in the city were not captured by the financial variables used in this study.

In general, the models fit to the full 43-year time series provided an adequate fit to the data. The full TSIR models (including waning of immunity) explained approximately 66% (range: 43-93%) of the variability in typhoid case counts over the study period (Table S3). Each financial predictor explained on average 47% of the variability in long-term typhoid transmission across the variables and cities (median city range: 8-63%; median financial predictor range: 2-90%) (Table S4).

We validated the models by fitting to the data through 1926 then using the fitted models to predict the last five years of typhoid mortality; the long-term typhoid transmission rate was predicted from water supply receipts. In most cases, the overall trend and seasonal peaks were captured, but the model projections could not explain some of the mortality spikes (Figures S12-S15). The models were generally a good fit for the data, with small out-of-sample mean squared prediction errors (Table S3).

Our results were not sensitive to methods of handling missing data and zeros. Seasonal transmission patterns remained the same, and long-term trends retained their general shape, but the intercepts varied depending on adjustments to imputation for missing values. Our results were also not sensitive to the maximum duration of immunity. All cities had different patterns and functions of immunity decay, but the shapes of the seasonal and long-term transmission rates of typhoid were mostly preserved when the models were fit assuming the maximum duration of immunity (∼13 years) or no waning of immunity.

## DISCUSSION

The decline in typhoid mortality in the early 20^th^ century U.S. has been attributed to investments in water and sewer systems. Our analysis solidifies this hypothesis. Furthermore, we characterized seasonal and long-terms trends in typhoid transmission and quantified the relationship between infrastructure investments and declines in transmission rates.

Historically, typhoid fever cases peaked during late summer/early fall in the U.S. (6, 27). Yearly peaks of typhoid transmission coincide with warmer temperatures, similar to global trends (7, 28–30). This pattern may be related to the enhanced growth of the bacteria at warmer temperatures, seasonal changes in diet (i.e. increased consumption of uncooked fruit and vegetables in summer and fall), or the increased abundance of flies that may serve as mechanical vectors of the bacteria (6, 7, 31, 32). Additional fluctuations in typhoid transmission seen in some cities might be explained by seasonal variation in rainfall, which typically peaks in spring and summer in the eastern U.S. and winter on the west coast, and can impact the water supply to a city (29, 30, 33).

We also found that seasonality in typhoid transmission varied by water source, which has a number of possible explanations. Cities relying on the Great Lakes for water had the least seasonal variability in transmission. Large bodies of water tend to be less impacted by seasonal changes in temperature and rainfall (34, 35). The Great Lakes have a moderating effect on climate, absorbing heat and cooling the air in the summer, yet radiating heat and protecting from frost in the fall (36, 37). Flowing water can slow down the movement of microbes (38), which may explain the lower seasonal variability in typhoid transmission among cities that draw water from rivers. Reservoirs and lakes are mostly smaller stagnant water sources, and may be more sensitive to seasonal changes in climate.

Differences in water source type may help to explain why some nearby cities exhibited different seasonal patterns. For example, New York and Philadelphia, though less than 100 miles apart, had different patterns of seasonal typhoid mortality and transmission (Figure 1, Figure S2). From 1890-1910, the typhoid mortality rate in New York was considerably lower than in Philadelphia (22.4 versus 43.1 deaths per 100,000 per year, respectively). However, typhoid transmission was more seasonal in New York (which relied on rural reservoirs) compared to Philadelphia (which drew its water from rivers). While typhoid transmission consistently peaked in the late summer/early fall in New York, Philadelphia also had smaller peaks in the spring and late fall. It is possible that these differences reflect differences in the predominant route of typhoid transmission (i.e. food-versus water-borne) in the two cities. Strong seasonality in typhoid incidence was also noted in Santiago, Chile in the 1970-80s, and was linked to seasonal irrigation of crops with contaminated wastewater; typhoid incidence declined sharply once this practice was ended (5, 39, 40). A better understanding of the drivers underlying seasonal patterns of typhoid transmission, and the differences noted among the various water sources, can aid typhoid control efforts.

In general, we found that investments in sewer systems were not as strongly associated with trends in typhoid transmission as water supply variables, most likely due to the fact that modern sewage treatment did not occur until later in the century (12). Furthermore, investments in the continued operation and maintenance of water and sewer systems had a larger and more immediate impact on typhoid transmission compared to investments in acquisition or construction. This may reflect increases in the impact of clean water and sanitation technologies over time (12). Associations also varied across cities, perhaps reflecting differences in water source types, public versus private ownership of water supplies, and rates of migration and poverty in the different cities. On average, the financial variables individually explained approximately half of the variability in typhoid transmission, and in several cases, explained over 90% of the variability. This supports the theory that these variables alone can account for the decline in typhoid transmission. However, other factors may also be important.

A previous study by Cutler and Miller showed similar findings (12). They estimated that on average, filtration and chlorination reduced typhoid fever mortality by 25% from 1900 to 1936. They claimed that clean water technologies explained almost all of the decline in typhoid mortality, estimating that the cost of clean water technologies per person-year saved was $500 in 2003 ($666 in 2017), suggesting it was highly cost-effective. However, their analysis did not consider the complexities of typhoid transmission, such as chronic carriers, host immunity, and interactions between susceptible and infectious individuals, which may bias their results.

It has heretofore been difficult to evaluate the benefits of water and sanitation infrastructure investments compared to the deployment of new typhoid conjugate vaccines without data to quantify the costs and impact of the former (41). With the recent World Health Organization recommendation for typhoid conjugate vaccine use and pilot studies underway (42, 43), governments are looking to prioritize the allocation of resources to yield the greatest decrease in typhoid burden. While long-term investments in water and sanitation systems are associated with decreased typhoid transmission, they also have benefits that extend beyond typhoid. Nevertheless, future studies should focus on comparing the cost-effectiveness and budget impact of the two interventions, bearing in mind the context and feasibility of deployment.

This study had some limitations. The weekly mortality counts likely suffer from lack of sensitivity and specificity in the diagnosis of typhoid fever. Additionally, we implicitly account for case fatality rates in our analysis. These issues are unlikely to bias our results provided the under- or over-reporting of typhoid mortality was consistent over the study period. The cities chosen for our analysis were also limited by data availability. As a result, all of the cities were primarily in the north-eastern U.S. All cities also had missing data, which had to be imputed. Furthermore, the roles of chronic carriers and immunity to typhoid are not fully understood. Our inclusion of carriers in the model matches the natural history of typhoid, but we did not examine whether it was necessary to model carriers separately. Patterns in the decay of immunity to typhoid varied widely across cities. Nevertheless, transmission rate estimates were not sensitive to the way we modelled immunity to infection. Finally, due to high levels of correlation between the financial variables, we were not able to consider combinations of water supply and sewer system variables. Some of the overall decline in transmission may have been attributable to other interventions such as economic and nutritional gains, and behaviour-change campaigns targeting hand and food washing (12, 44–46).

Our results aid in the understanding of the dynamics of typhoid transmission and potential impact of improvements in water and sanitation infrastructure, which is still lacking in many parts of the world. Worldwide, approximately 1.1 billion people lack access to clean water, and roughly 2.5 billion people lack adequate sanitation (47). Water and sanitation technologies can have substantial health returns. However, resource-poor countries must prioritize spending on public health issues, bearing in mind the cost-effectiveness and affordability of interventions. Our results can help to inform comparative cost-effectiveness analyses of different interventions to reduce the global burden of typhoid fever.

## NOTES

### Authors’ contributions

VEP and BTG conceived of the study and designed the study. MTP carried out the data analyses. MTP and VEP drafted the manuscript. KAO participated in data analysis. All authors critically revised the manuscript and gave final approval for publication and agree to be held accountable for the work performed herein.

