## Supplemental Material for "Changes in historical typhoid transmission across 16 U.S. cities, 1889-1931: Quantifying the impact of investments in water and sewer infrastructures"

*Supplement S1: Model-fitting process.*

We fit Time-series Susceptible-Infectious-Recovered (TSIR) models [22] to each city’s time series to investigate seasonal and long-terms trends in typhoid transmission rates. TSIR models estimate the disease transmission rate using a regression framework by reconstructing the underlying susceptible and infectious populations. New infections at time *t*+1 (*I_t_*_+1_) arise from transmission from infectious (*I_t_*) to susceptible (*S_t_*) individuals at time *t*:

$I_{t+1}=\beta_{t}I_{t}^{\alpha}S_{t}$	*(1)*

where $\beta_{t}$ is the disease transmission rate at time *t* and $\alpha$ is a scaling factor that adjusts for heterogeneous mixing in the population.

We reconstructed the infectious ($I_{t}$) and susceptible ($S_{t}$) populations based on the typhoid mortality and census data using the first ten years of data (prior to the introduction of water and sanitation interventions in each city) as outlined by Bjørnstad et al [23, 24]. The susceptible population at time *t* is equal to the susceptible population at the previous time step plus the number of births minus the number of new infections, which can be summarized as follows:

$S_{t}=\bar{S}+D_{0}+\sum_{k=0}^{t-1} B_{k}-\sum_{k=0}^{t-1} I_{k}$	*(2)*

where $\bar{S}$ is the mean susceptible population size, $D_{0}$ is the deviation at time zero, $\sum_{k=0}^{t-1} B_{k}$ is the sum of all weekly births up to time *t*, and $\sum_{k=0}^{t} I_{k}$ is the sum of all infections up to time *t*. The number of “true” infections at time *t* is estimated from the observed deaths at time *t* divided by an underreporting factor *ρ* ($I_{t}=Y_{t}/\rho$). Equation 2 can be rearranged as:

$\sum_{k=0}^{t} B_{k}=\left( \frac{1}{\rho} \right)\sum_{k=0}^{t} Y_{k}-D_{0}+D_{t}$	*(3)*

to estimate the underreporting fraction (1/slope), deviation at time zero (*D*_0_, intercept), and model residuals ($D_{t}=S_{t}-\bar{S})$.

We modified Equation 1 to account for the unique features of typhoid epidemiology, including the contribution of chronic carriers (*C*) to the overall prevalence of infection. Furthermore, we separated the transmission parameter $\beta_{t}$ into seasonal and long-term components ($\beta_{seas}$ and $\beta_{lt}$, respectively). Thus, the TSIR model for typhoid is as follows:

$I_{t+1}=\beta_{lt}\beta_{seas}{(I_{t}+C)}^{\alpha}S_{t}$	*(4)*

Equation 4 is then log-transformed to obtain an additive model:

$log{(I}_{t+1})={log(\beta}_{lt})+\log\left( \beta_{seas} \right)+\alpha\log\left( I_{t}+C \right)+log(S_{t})$	*(5)*

To estimate *C*, we maximized the likelihood of the fitted regression over values ranging from 0 to *N* (the population size).

To estimate the parameters of our main model, we first fit the parametric part (everything except for the long-term transmission parameter) of Equation 5 using weighted least-squares regression:

$log{(I}_{t+1})=\log\left( \beta_{seas} \right)+\alpha\log\left( I_{t}+CC \right)+log(S_{t})$	*(6)*

The susceptible population at time *t* can be expressed as a function of the total population at time *t* minus the previously infectious and recovered individuals: $S_{t}=N_{t}-\sum_{i=0}^{m} I_{t-i}\kappa_{i}$, where $\kappa_{i}$is the degree of immunity *i* generation intervals after infection (determined by a decay of immunity function). Using this equation and a logarithmic transformation and first-degree Taylor series expansion around the average susceptible population size, $\log\left( S_{t} \right)$can be initially approximated using the previous equation: $\log\left( S_{t} \right)\approx\left[ \log\left( S_{mean} \right)+\frac{N_{t}}{S_{mean}}-1 \right]-\sum_{i=0}^{m} I_{t-i}\kappa_{i}.$ Using the approximations of $log(S_{t})$, this equation can be rewritten:

$log{(I}_{t+1})=\log\left( \beta_{seas} \right)+\alpha\log\left( I_{t}+C \right)-\sum_{i=0}^{m} I_{t-i}\kappa_{i}+\left[ \log\left( S_{mean} \right)+\frac{N_{t}}{S_{mean}}-1 \right]$	*(7)*

We used the value of *C* estimated from Equation 5 above $(log{(I}_{t+1})={log(\beta}_{lt})+\log\left( \beta_{seas} \right)+\alpha\log\left( I_{t}+C \right)+log(S_{t}))$, and estimated 13 values of $\beta_{seas}$ corresponding to transmission during the same four-week period each year. We then used the semi-parametric method described by Koelle and Pascual [1, 2] to estimate the variation in $\beta_{lt}$ over the full 43-year study period from 1889-1931. To fit the model, we used a back-fitting algorithm comprised of repeated penalized cubic splines on $\kappa$, recursive first-order Taylor series expansion approximations on the $log(S_{t})$ term, and weighted least squares regressions on iterations of Equation 7. The iterative process was necessary to allow for the more accurate approximation of the Taylor series expansion and the cubic splines to converge. For the weighted least squares regressions, the weights were calculated as **I-W**, where **I** was the identity matrix of the same dimension, and **W** was the truncated Gaussian kernel weight matrix calculated from the spline penalty weights.

The penalized cubic splines on $\kappa$ account for non-linearity in the duration and decay patterns in immunity. Not much is known about the duration of immunity to typhoid, so a sensitivity analysis was also performed to determine the maximum duration of immunity to use for this model.

After the above algorithm converged, we used the same truncated Gaussian kernel weight matrix **W** to smooth the residuals of the model fit with all other parameters estimated. The smoothed residuals were then used to estimate the nonparametric variation in the long-term transmission rate ($\beta_{lt}$). The final long-term transmission rate was then multiplied by the population at each time point to create per capita estimates.

The smoothing parameter and spline penalty weights were selected using cross-validation and testing across a range of values. Each city was fit using all possible values of the smoothing parameter from one to 35 in one-unit increments and spline penalty values from negative two to 30 in one-unit increments. If the parameters chosen were at the ends of the respective ranges, the intervals were extended until the optimal values fell within the values tested. To identify the optimal model, we used leave-one-out validation, in which we dropped one data point at a time (for all data points), fit the model to the remaining data, and then used the fitted model to predict the out-of-sample data point. We calculated the sum of squared differences between each point and its out-of-sample prediction over all points, and stored this as the cross-validated (CV) value for each model fit. The optimal model for each city was the one with the smallest CV value.

To validate the TSIR models and assess their predictive ability, we went back and fit each TSIR model to the first 38 years of data (1889-1926). Using the fitted model parameters, we projected forward for the last five years (1927-1931) and compared the observed and predicted typhoid mortality. To predict the long-term typhoid transmission rate, we used water supply receipts, since this variable provided the best fit for most cities, and we wanted to use the same variable for all cities to be consistent.

***Supplemental Figures***

**Figure S1. Map of 16 cities, with water supply types.** Each city included in the analysis is denoted by a different colour in its geographical location in the United States. Squares denote cities with reservoirs, triangles denote those using the Great Lakes, and circles denote those with rivers as their main water source.

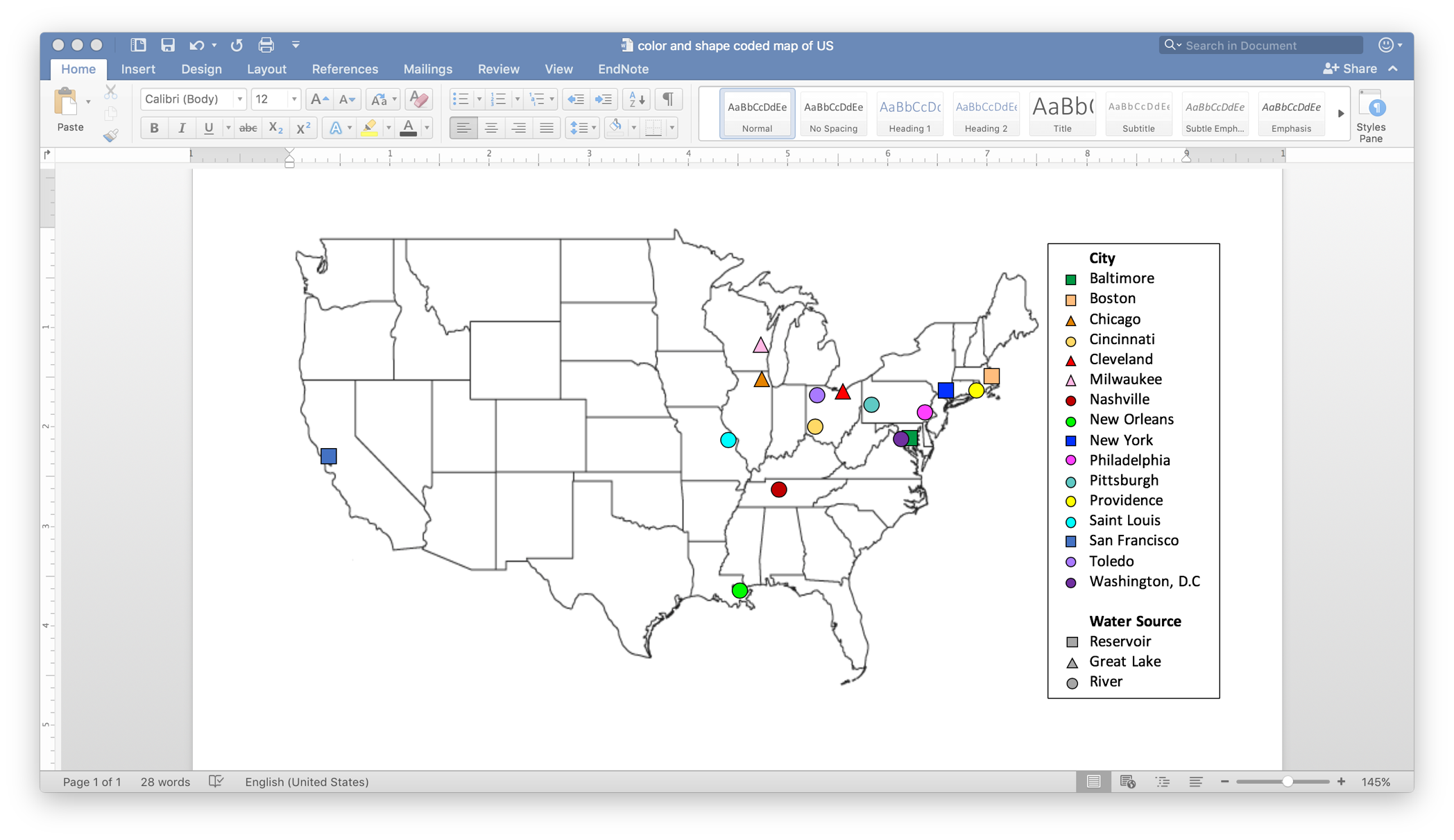

**Figure S2. Weekly time-series of reported typhoid mortality in each city.** The observed time series of weekly deaths reportedly due to typhoid (black lines) and the yearly typhoid deaths per 100,000 people (red Xs) is shown for each city from 1889-1931.

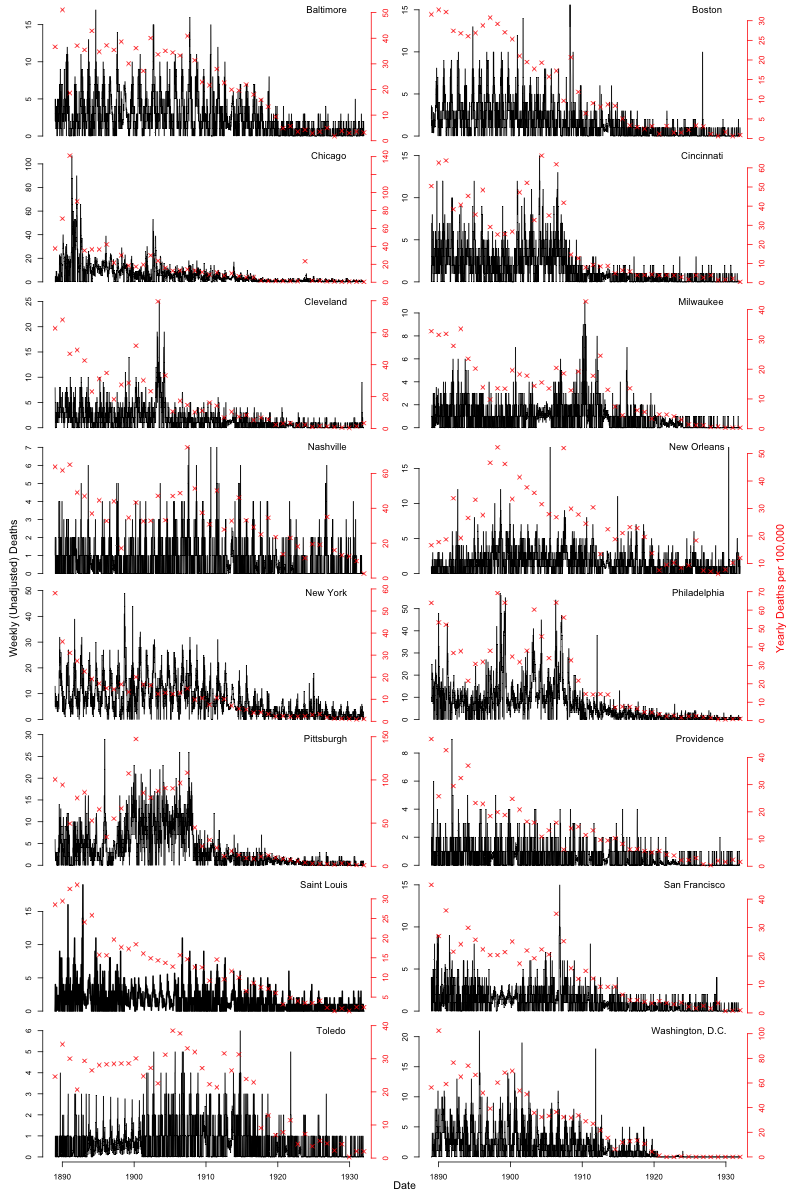

**Figure S3. Seasonal transmission: Simple TSIR models for pre- and post- water supply intervention.** The estimated four-week seasonal transmission rates extracted from each city’s simple TSIR model are shown for each pre- (red) and post- (blue) water supply intervention period.

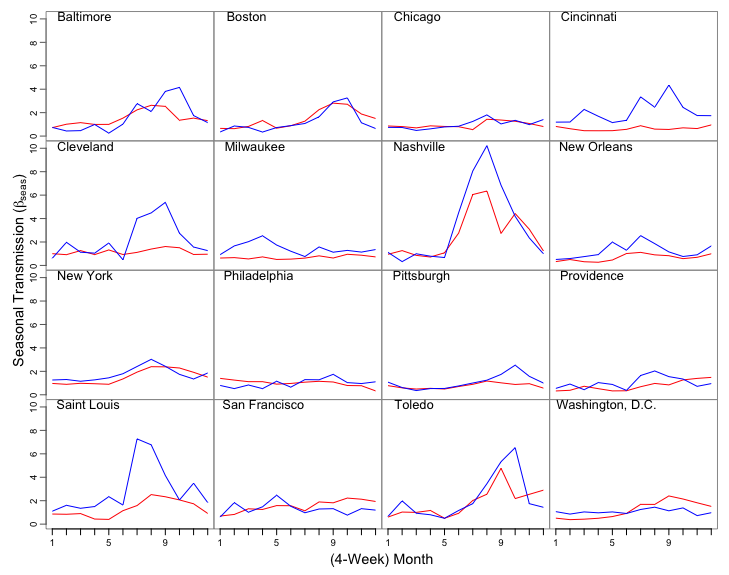

### ***Figures S4-11. Financial variable time series.***

The yearly time series of each of the eight financial water supply or sewer system variables is plotted over the study period in per capita increments. Dollar amounts are not adjusted for inflation.

**Figure S4. Per Capita Water Supply Receipts.** Annual water supply receipts from 1902-1931 are shown for each city in per capita increments (US$ per person).

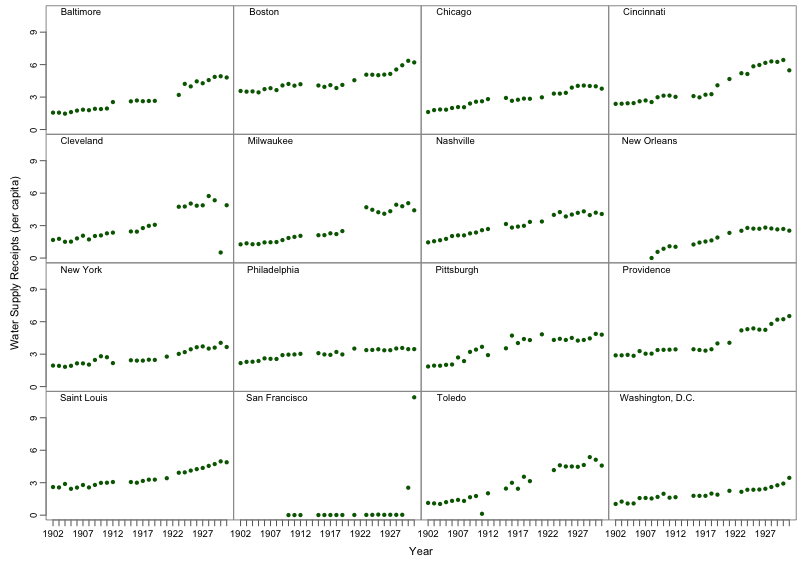

#### *Figure S5. Financial variable time series: Per Capita Water Supply Expenses.*

Annual spending on water supply expenses from 1902-1931 is shown for each city in per capita increments.

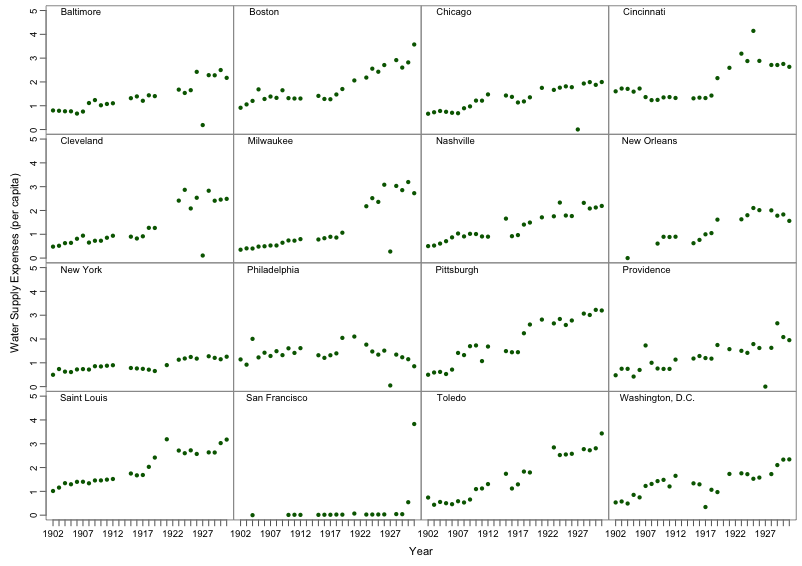

#### *Figure S6. Financial variable time series: Per Capita Sewer System Expenses.*

Annual spending on sewer system expenses from 1902-1931 is shown for each city in per capita increments.

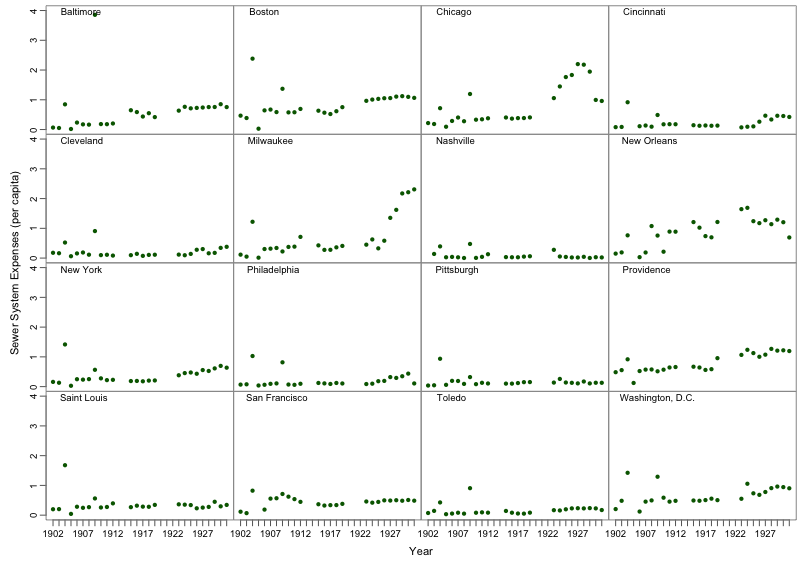

#### *Figure S7. Financial variable time series: Per Capita Water Supply Outlays.*

The per-capita water supply outlays estimate is plotted in green dots for each city and time point, while the year in which interventions were introduced are represented by the dashed lines for filtration (red), chlorination (blue), or other interventions (purple). Outlier not seen: in 1930, water supply outlays from San Francisco totalled $70.97 per capita in water supply receipts. Also not seen: Chicago and New Orleans introduced water supply interventions in 1900, prior to the time period shown.

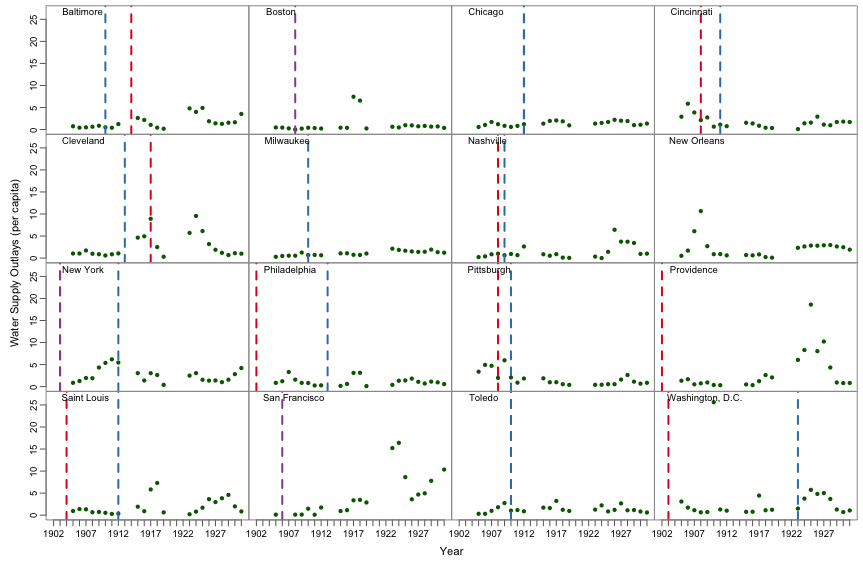

#### *Figure S8. Financial variable time series: Per Capita Sewer System Outlays.*

Annual spending on sewer system outlays from 1902-1931 is shown for each city in per capita increments.

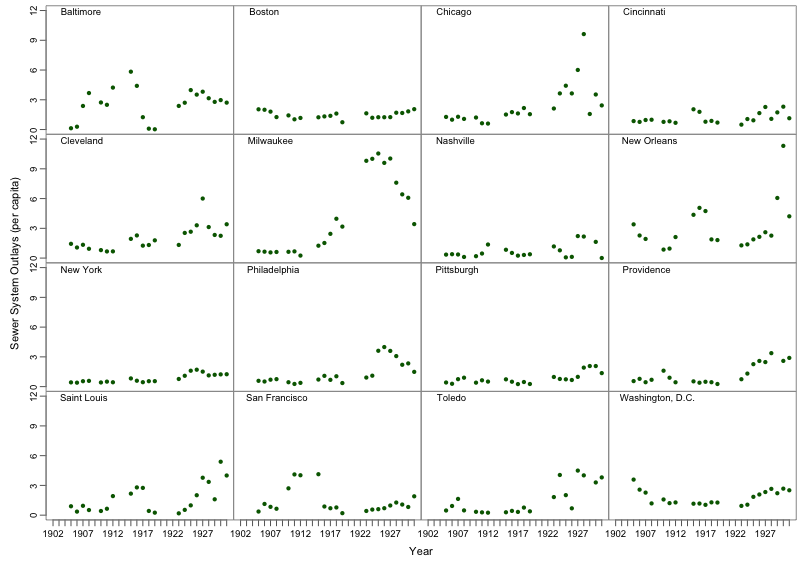

#### *Figure S9. Financial variable time series: Per Capita Value of the Water Supply System.*

The overall annual value of the water supply system from 1902-1931 is shown for each city in per capita increments. Outliers not seen: in 1897, it totalled 508.55$ per capita in Washington, D.C.

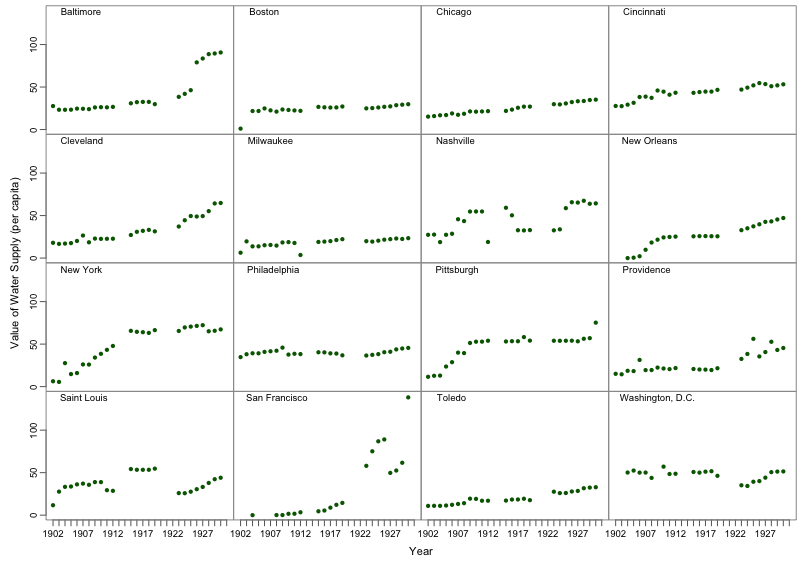

#### *Figure S10. Financial variable time series: Per Capita Funded Debt of the Water System.*

The overall annual accrued debt and/or funded loans for the water supply system from 1902-1931 is shown for each city in per capita increments. Note: Data were not available for this variable in Washington, D.C.

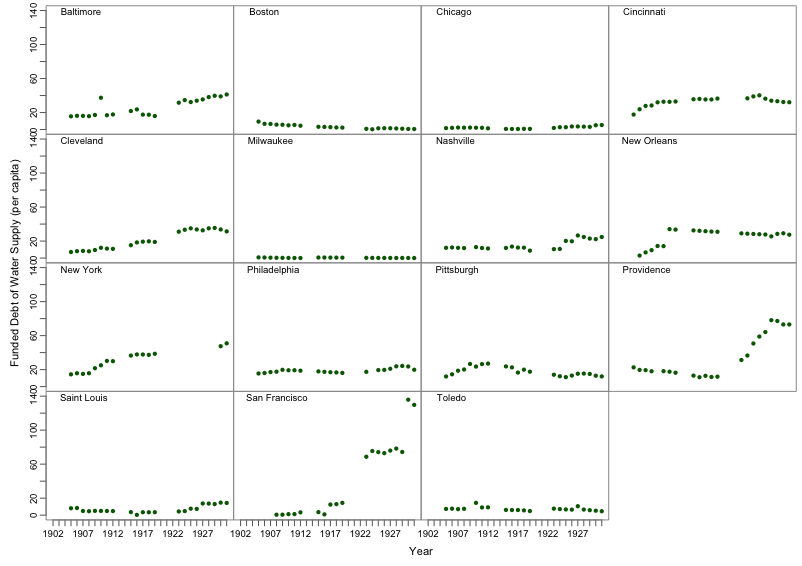

#### *Figure S11. Financial variable time series: Per Capita Funded Debt of the Sewer System.*

The overall annual accrued debt and/or funded loans for the sewer system from 1902-1931 are shown for each city in per capita increments. Note: Data were not available for this variable in Washington, D.C.

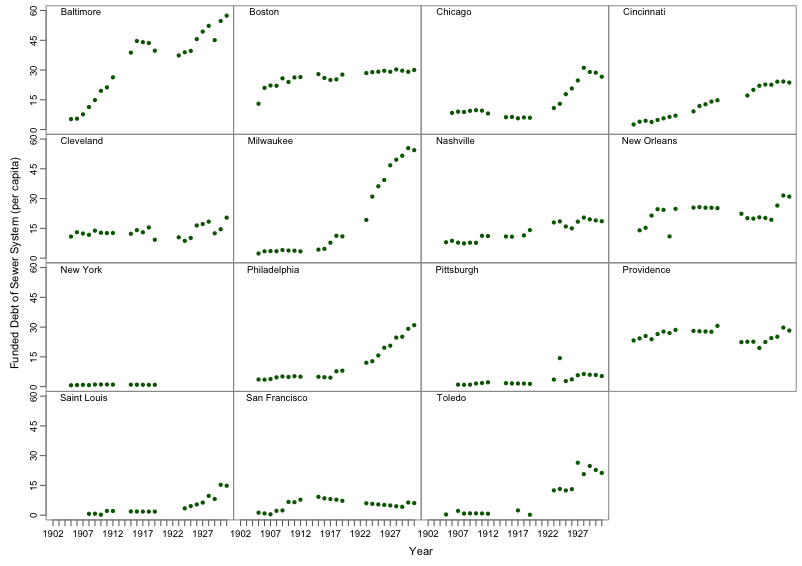

### ***Figures S12-S15. TSIR predictions.***

For each city, the TSIR model is fit using the first 38 years of data, then used to predict the last 5 years of data.

#### *Figure S12. TSIR predictions for Baltimore, Boston, Chicago, and Cincinnati.*

In each plot, the observed data is shown in black, the model fit to the first 38 years is shown in blue, and the predicted last 5 years is shown in red.

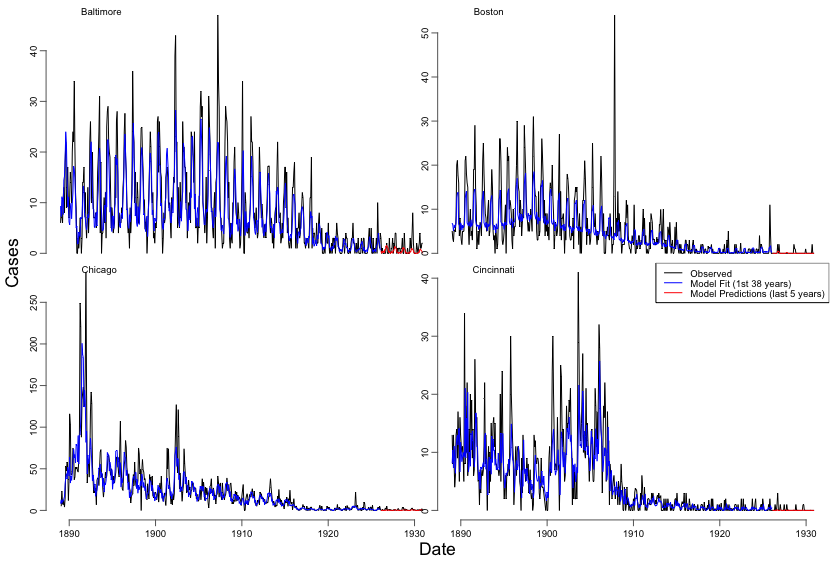

#### *Figure S13. TSIR predictions for Cleveland, Milwaukee, Nashville, and New Orleans.*

In each plot, the observed data is shown in black, the model fit to the first 38 years is shown in blue, and the predicted last 5 years is shown in red.

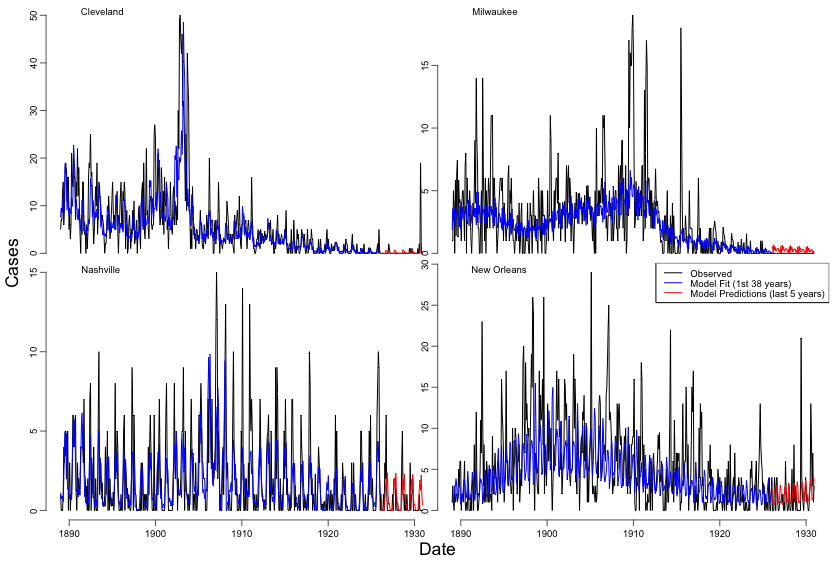

#### *Figure S14. TSIR predictions for New York, Philadelphia, Pittsburgh, and Providence.*

In each plot, the observed data is shown in black, the model fit to the first 38 years is shown in blue, and the predicted last 5 years is shown in red.

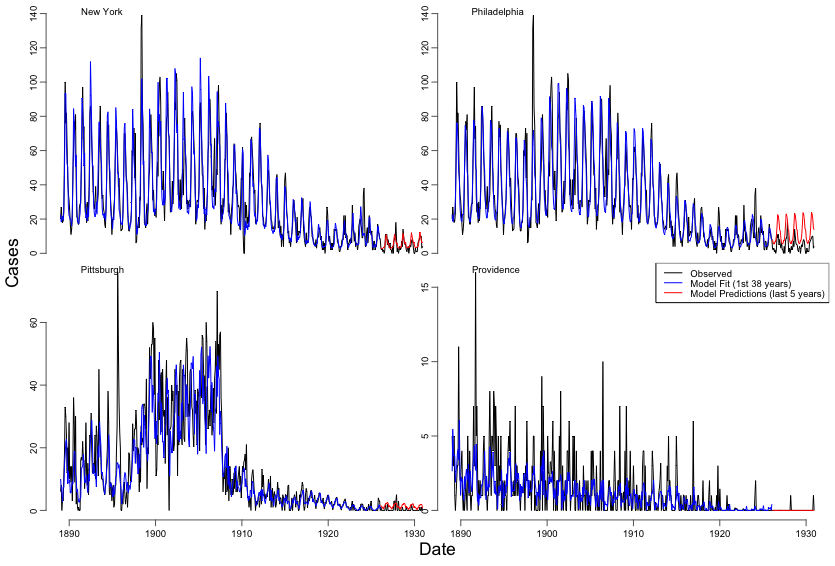

#### *Figure S15. TSIR predictions for St. Louis, San Francisco, Toledo, and Washington, D.C.*

In each plot, the observed data is shown in black, the model fit to the first 38 years is shown in blue, and the predicted last 5 years is shown in red.

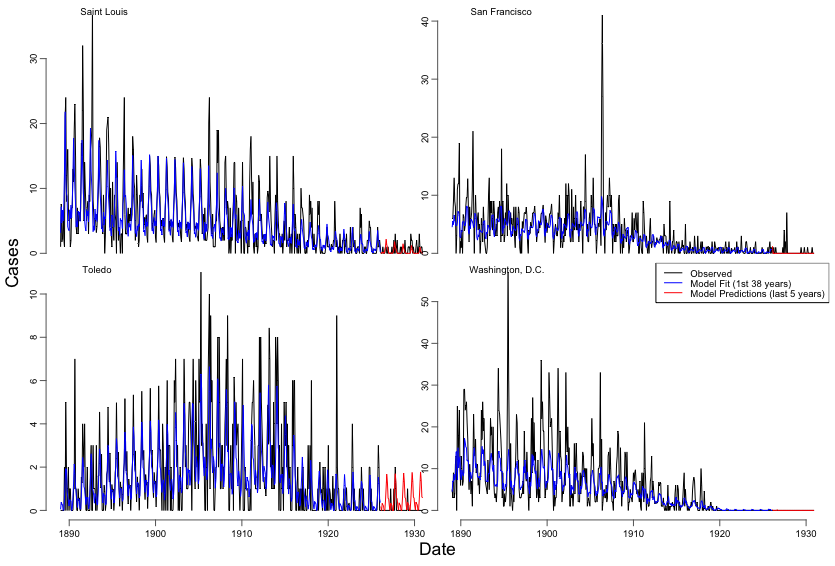

### ***Table S1. References for water supply interventions and dates.***

All references used for data on water supply interventions and water source types were extracted from a variety of sources, noted below. Most cities had data available from the U.S. Census Bureau in addition to individual municipal sources, noted in the table as “U.S. Census Bureau (Yes/No)”.

| **City** | **State** | **Water Source** | **U.S. Census Bureau (Yes/No)** | **Additional Sources** |
| --- | --- | --- | --- | --- |
| San Francisco | CA | Arroyo de la Laguna, Alameda Creek, artesian wells in Pleasanton (Hetch Hetchy reservoir in Yosemite since 1923); owned/operated by private company -- Spring Valley Water Co | No | <http://www.sfmuseum.org/hist3/perry.html> |
| Washington, D.C. | - | Potomac River via Washington Aqueduct; McMillan Reservoir and Bryant Street high-lift pump after 1905 | Yes (Filtration, Chlorination) | <https://www.dcwater.com/history-water-system> |
| Chicago | IL | Lake Michigan; flow direction of Chicago River reversed in 1900 to prevent waste water from entering lake | Yes (Filtration, Chlorination) | <http://encyclopedia.chicagohistory.org/pages/1325.html> |
| New Orleans | LA | Rainwater cisterns, Mississippi River, artesian wells; city subject to regular flooding at late as mid-1880s; drainage plan initiated in 1896; water treatment authorized/built in 1899 | Yes (Filtration) | <http://www.swbno.org/history_history.asp> |
| Boston | MA | Long Pond (Lake Cochituate) via Cochituate aqueduct and Brookline Reservoir; several reservoirs, aqueducts built between 1864 and 1900; Wachusett Reservoir/Dam/aqueduct completed in 1908 | Yes (Filtration) | <http://www.bwsc.org/ABOUT_BWSC/systems/water/Water_history.asp> |
| Baltimore | MD | Jones Falls reservoir, Lake Roland/Lake Hampden/Mount Royal reservoirs (1862), Druid Hill reservoir (1873), Loch Raven reservoir and Lake Montebello (1881); chlorination 1910, filtration 1915 | Yes (Filtration, Chlorination) | <http://cityservices.baltimorecity.gov/dpw/waterwastewater02/waterquality3.html> <http://www.baltimorecity.gov/Government/AgenciesDepartments/PublicWorks/BureauofWaterWastewater/FactSheet.aspx> |
| Saint Louis | MO | Mississippi River (via High Service pumping station at Bissels Point since 1871, also via Low Service Chain of Rocks plant since 1894); filtration plant dedicated in 1915 | Yes (Filtration, Chlorination) | <http://www.stlwater.com/history2.php> |
| New York | NY | Old Croton Reservoir (via Old Croton aqueduct, since 1842); distribution reservoirs at 42nd St, Central Park, Boyds Corner, Middle Branch; additional reservoirs in Catskills in 1905-1915 | Yes (Filtration, Chlorination) | <http://www.nyc.gov/html/dep/html/drinking_water/history.shtml> |
| Cincinnati | OH | Ohio River (with subsiding, storage, & filtering reservoirs, e.g. Eden Park) | Yes (Filtration, Chlorination) | <http://books.google.com/books?id=6vzVAAAAMAAJ&pg=PA42&lpg=PA42&dq=cincinnati+water+works+history&source=bl&ots=Qe82LaTBvG&sig=zGhGv6VH7yZ2LwXvdLySJ7B8OnI&hl=en&sa=X&ei=um7rTuCUPIT30gGDgpXQCQ&ved=0CGIQ6AEwBw#v=onepage&q=cincinnati%20water%20works%20history&f=false> |
| Cleveland | OH | Lake Erie west of the Cuyahoga River; off-shore intake (Kinsman Reservoir) began operation in 1885; further off-shore intakes created 1890-1916; chlorination began in 1911, daily testing 1913, filtration 1917 | Yes (Filtration, Chlorination) | <http://www.clevelandwater.com/about_us/history.aspx> |
| Toledo | OH | Maumee River; filtration plant upstream of Broadway Pumping Station began operation in 1910; source changed to Lake Erie in 1940s | Yes (Filtration, Chlorination) | <http://www.ci.toledo.oh.us/Departments/PublicUtilities/DivisionofWaterTreatment/HistoryofWaterTreatmentinToledo/tabid/375/Default.aspx> |
| Philadelphia | PA | Schuylkill River, Delaware River, Leigh River, various creeks | Yes (Filtration, Chlorination) | [http://www.phillyh2o.org/ http://www.phila.gov/water/PWD_Historical.html](http://www.phillyh2o.org/) |
| Pittsburgh | PA | Allegheny River (with holding reservoirs); filtration began in 1909 (Southside) and 1914 (Northside); chlorination began in 1911; City of Pittsburgh merged with City of Allegheny (Northside) in 1907 | Yes (Filtration, Chlorination) | <http://www.pgh2o.com/history02.htm> |
| Providence | RI | Pawtuxet River (at Cranston); filtration began 1906; filtered water stored in 3 open reservoirs; shortages during dry weather; Scituate Reservoir and treatment plant constructed in 1926 (by damming river) | Yes (Filtration) | <http://www.provwater.com/history.htm> |
| Nashville | TN | Cumberland River via reservoirs used for settling and storage; pumping station relocated upstream of Brown's Creek in 1889; treatment with hypochlorite in 1908, liquid chlorine in 1920, filtration in 1928 | Yes (Filtration, Chlorination) | <http://www.nashville.gov/water/docs/other/water_history_additional_information.pdf> |
| Milwaukee | WI | Lake Michigan via Kilbourne Reservoir; second pumping station at Milwaukee River in 1924 | Yes (Chlorination) | <http://city.milwaukee.gov/water/customer/FAQs/additionalinfo#4> |

### ***Table S2. Harmonic regression estimates and seasonal amplitudes of typhoid mortality: pre- and post- water supply intervention in each city.***

Time trends and seasonal amplitudes were estimated for each city pre- and post- intervention in preliminary analyses with harmonic regression. Values shown in grey were not statistically significant at the 0.05-level, while values in black had p-values<0.05. In the last column, the ratio (post-/pre- water supply intervention) was calculated from the six-month and one-year amplitudes estimated from the regression models.

|  | **Pre-Intervention** | | | | **Post-Intervention** | | | | | *Ratio (Post/Pre)*  *of Seasonal Amplitude* | |
| --- | --- | --- | --- | --- | --- | --- | --- | --- | --- | --- | --- |
|  | *Time trend* | | *Seasonal Amplitude* | | *Time trend* | | *Seasonal Amplitude* | | |  |  |
|  | *intercept* | *slope* | *6-mo.* | *1-yr.* | *intercept* | *slope* | *6-mo.* | | *1-yr.* | *6-mo.* | *1-yr.* |
| **Baltimore** | -3.35 | 0.00 | 0.15 | 0.62 | 76.94 | -0.04 | 0.08 | 0.24 | | 0.49 | 0.39 |
| **Boston** | 27.10 | -0.01 | 0.22 | 0.55 | 25.75 | -0.01 | 0.04 | 0.07 | | 0.19 | 0.13 |
| **Chicago** | 27.78 | -0.01 | 0.14 | 0.26 | 141.03 | -0.07 | 0.04 | 0.18 | | 0.32 | 0.68 |
| **Cincinnati** | -18.04 | 0.01 | 0.15 | 0.18 | 4.85 | -0.00 | 0.00 | 0.02 | | 0.03 | 0.10 |
| **Cleveland** | 41.50 | -0.02 | 0.15 | 0.08 | 17.29 | -0.01 | 0.04 | 0.06 | | 0.25 | 0.67 |
| **Milwaukee** | 2.28 | 0.00 | 0.02 | 0.07 | 27.95 | -0.01 | 0.04 | 0.06 | | 1.91 | 0.81 |
| **Nashville** | -11.36 | 0.01 | 0.06 | 0.17 | 12.44 | -0.01 | 0.07 | 0.14 | | 1.16 | 0.83 |
| **New Orleans** | -151.79 | 0.08 | 0.14 | 0.13 | 43.97 | -0.02 | 0.14 | 0.20 | | 0.97 | 1.51 |
| **New York** | -23.70 | 0.01 | 0.12 | 0.74 | 162.40 | -0.08 | 0.11 | 0.47 | | 0.88 | 0.64 |
| **Philadelphia** | 27.96 | -0.01 | 0.17 | 0.16 | 199.33 | -0.10 | 0.05 | 0.04 | | 0.32 | 0.24 |
| **Pittsburgh** | -154.97 | 0.08 | 0.14 | 0.23 | 64.03 | -0.03 | 0.04 | 0.11 | | 0.31 | 0.49 |
| **Providence** | 12.81 | -0.01 | 0.03 | 0.12 | 7.48 | -0.00 | 0.01 | 0.03 | | 0.43 | 0.26 |
| **Saint Louis** | 31.39 | -0.02 | 0.21 | 0.47 | 43.31 | -0.02 | 0.09 | 0.24 | | 0.43 | 0.51 |
| **San Francisco** | 16.32 | -0.01 | 0.08 | 0.21 | 20.84 | -0.01 | 0.02 | 0.04 | | 0.26 | 0.18 |
| **Toledo** | -25.43 | 0.01 | 0.06 | 0.12 | 26.07 | -0.01 | 0.03 | 0.08 | | 0.50 | 0.66 |
| **Washington, D.C.** | 15.94 | -0.01 | 0.07 | 0.64 | 53.81 | -0.03 | 0.05 | 0.17 | | 0.77 | 0.26 |

### ***Table S3. Variability in typhoid cases explained (R^2^) by full TSIR models, within-sample mean squared errors (MSE) and out-of-sample mean squared prediction errors (MSPE) from prediction models, by city.***

R^2^ is shown for the models fit to the full study period (1889-1931). MSE is shown for 1922-1926, the last five years of within-sample data used to predict the out-of-sample estimates. MSPE is shown for 1927-1931, the out-of-sample estimates predicted.

| **City** | **R^2^ (full model fit)** | **MSE (within-sample)** | **MSPE (out-of-sample)** | **Ratio (MSPE/MSE)** |
| --- | --- | --- | --- | --- |
| **Baltimore** | 0.65 | 2.90 | 2.61 | 0.90 |
| **Boston** | 0.66 | 2.10 | 0.25 | 0.12 |
| **Chicago** | 0.87 | 8.56 | 1.36 | 0.16 |
| **Cincinnati** | 0.77 | 0.41 | 0.26 | 0.63 |
| **Cleveland** | 0.72 | 0.82 | 5.54 | 6.76 |
| **Milwaukee** | 0.54 | 0.19 | 0.11 | 0.58 |
| **Nashville** | 0.44 | 2.38 | 1.33 | 0.56 |
| **New Orleans** | 0.53 | 6.64 | 10.29 | 1.55 |
| **New York** | 0.93 | 39.11 | 7.86 | 0.20 |
| **Philadelphia** | 0.86 | 39.03 | 75.53 | 1.94 |
| **Pittsburgh** | 0.83 | 0.63 | 1.67 | 2.65 |
| **Providence** | 0.49 | 0.14 | 0.03 | 0.21 |
| **Saint Louis** | 0.59 | 1.82 | 0.88 | 0.48 |
| **San Francisco** | 0.62 | 0.40 | 1.17 | 2.93 |
| **Toledo** | 0.43 | 0.42 | 0.49 | 1.17 |
| **Washington, D.C.** | 0.70 | 0.04 | 0.00 | 0.00 |

### ***Table S4. Variability in long-term typhoid transmission explained by financial water supply and sewer system variables, by city.***

*Values shown are the R^2^ numbers from the linear regression analyses for each city and financial variable.*

|  | **Baltimore** | **Boston** | **Chicago** | **Cincinnati** | **Cleveland** | **Milwaukee** | **Nashville** | **New Orleans** | **New York** | **Philadelphia** | **Pittsburgh** | **Providence** | **St. Louis** | **San Francisco** | **Toledo** | **Washington, D.C.** | **minimum** | **median** | **maximum** |
| --- | --- | --- | --- | --- | --- | --- | --- | --- | --- | --- | --- | --- | --- | --- | --- | --- | --- | --- | --- |
| **Water supply receipts** | 0.91 | 0.66 | 0.85 | 0.62 | 0.53 | 0.85 | 0.75 | 0.97 | 0.82 | 0.78 | 0.85 | 0.69 | 0.90 | 0.08 | 0.90 | 0.74 | *0.08* | ***0.80*** | *0.97* |
| **Water supply expenses** | 0.52 | 0.59 | 0.44 | 0.29 | 0.52 | 0.64 | 0.70 | 0.87 | 0.64 | 0.03 | 0.86 | 0.39 | 0.90 | 0.07 | 0.90 | 0.46 | *0.03* | ***0.55*** | *0.90* |
| **Sewer system expenses** | 0.02 | 0.04 | 0.46 | 0.00 | 0.01 | 0.35 | 0.03 | 0.57 | 0.08 | 0.00 | 0.05 | 0.64 | 0.02 | 0.00 | 0.00 | 0.09 | *0.00* | ***0.04*** | *0.64* |
| **Water supply outlays** | 0.36 | 0.05 | 0.24 | 0.49 | 0.10 | 0.61 | 0.09 | 0.03 | 0.06 | 0.01 | 0.49 | 0.30 | 0.13 | 0.17 | 0.00 | 0.01 | *0.00* | ***0.11*** | *0.61* |
| **Sewer system outlays** | 0.05 | 0.02 | 0.38 | 0.13 | 0.46 | 0.72 | 0.14 | 0.08 | 0.60 | 0.48 | 0.36 | 0.34 | 0.28 | 0.03 | 0.50 | 0.00 | *0.00* | ***0.31*** | *0.72* |
| **Value water supply system** | 0.66 | 0.45 | 0.93 | 0.85 | 0.75 | 0.43 | 0.12 | 0.86 | 0.81 | 0.11 | 0.68 | 0.57 | 0.03 | 0.51 | 0.84 | 0.07 | *0.03* | ***0.61*** | *0.93* |
| **Funded debt water supply** | 0.64 | 0.93 | 0.20 | 0.74 | 0.91 | 0.13 | 0.41 | 0.45 | 0.91 | 0.41 | 0.19 | 0.45 | 0.36 | 0.56 | 0.13 | - | *0.13* | ***0.45*** | *0.93* |
| **Funded debt sewer system** | 0.77 | 0.69 | 0.44 | 0.72 | 0.07 | 0.73 | 0.91 | 0.15 | 0.07 | 0.80 | 0.42 | 0.04 | 0.63 | 0.39 | 0.75 | - | *0.04* | ***0.63*** | *0.91* |
| **minimum** | *0.02* | *0.02* | *0.20* | *0.00* | *0.01* | *0.13* | *0.03* | *0.03* | *0.06* | *0.00* | *0.05* | *0.04* | *0.02* | *0.00* | *0.00* | *0.00* | *0.00* | *0.02* | *0.20* |
| **median** | ***0.58*** | ***0.52*** | ***0.44*** | ***0.55*** | ***0.49*** | ***0.63*** | ***0.28*** | ***0.51*** | ***0.62*** | ***0.26*** | ***0.45*** | ***0.42*** | ***0.32*** | ***0.13*** | ***0.63*** | ***0.08*** | *0.08* | ***0.47*** | *0.63* |
| **maximum** | *0.91* | *0.93* | *0.93* | *0.85* | *0.91* | *0.85* | *0.91* | *0.97* | *0.91* | *0.80* | *0.86* | *0.69* | *0.90* | *0.56* | *0.90* | *0.74* | *0.56* | *0.90* | *0.97* |

***Supplement references.***

1. Koelle K, Pascual M. Disentangling extrinsic from intrinsic factors in disease dynamics: a nonlinear time series approach with an application to cholera. Am Nat **2004**; 163:901-13.

2. Koelle K, Rodó X, Pascual M, Yunus M, Mostafa G. Refractory periods and climate forcing in cholera dynamics. Nature **2005**; 436:696.
